# Novel, provable algorithms for efficient ensemble-based computational protein design and their application to the redesign of the c-Raf-RBD:KRas protein-protein interface

**DOI:** 10.1101/790949

**Authors:** Anna U. Lowegard, Marcel S. Frenkel, Jonathan D. Jou, Adegoke A. Ojewole, Graham T. Holt, Bruce R. Donald

**Affiliations:** Program in Computational Biology and Bioinformatics, Duke University Medical Center, Durham, NC, USA; Department of Computer Science, Duke University, Durham, NC, USA; Department of Biochemistry, Duke University Medical Center, Durham, NC, USA

## Abstract

The *K*\* algorithm provably approximates partition functions for a set of states (e.g., protein, ligand, and protein-ligand complex) to a user-specified accuracy *ε*. Often, reaching an *ε*-approximation for a particular set of partition functions takes a prohibitive amount of time and space. To alleviate some of this cost, we introduce two algorithms into the osprey suite for protein design: fries, a Fast Removal of Inadequately Energied Sequences, and *EWAK*\*, an Energy Window Approximation to *K*\*. In combination, these algorithms provably retain calculational accuracy while limiting the input sequence space and the conformations included in each partition function calculation to only the most energetically favorable. This combined approach leads to significant speed-ups compared to the previous state-of-the-art multi-sequence algorithm, *BBK*\*. As a proof of concept, we used these new algorithms to redesign the protein-protein interface (PPI) of the c-Raf-RBD:KRas complex. The Ras-binding domain of the protein kinase c-Raf (c-Raf-RBD) is the tightest known binder of KRas, a historically “undruggable” protein implicated in difficult-to-treat cancers including pancreatic ductal adenocarcinoma (PDAC). fries/*EWAK*\* accurately retrospectively predicted the effect of 38 out of 41 different sets of mutations in the PPI of the c-Raf-RBD:KRas complex. Notably, these mutations include mutations whose effect had previously been incorrectly predicted using other computational methods. Next, we used fries/*EWAK*\* for prospective design and discovered a novel point mutation that improves binding of c-Raf-RBD to KRas in its active, GTP-bound state (KRas^GTP^). We combined this new mutation with two previously reported mutations (which were also highly-ranked by osprey) to create a new variant of c-Raf-RBD, c-Raf-RBD(RKY). fries/*EWAK*\* in osprey computationally predicted that this new variant would bind even more tightly than the previous best-binding variant, c-Raf-RBD(RK). We measured the binding affinity of c-Raf-RBD(RKY) using a bio-layer interferometry (BLI) assay and found that this new variant exhibits single-digit nanomolar affinity for KRas^GTP^, confirming the computational predictions made with fries/*EWAK*\*. This study steps through the advancement and development of computational protein design by presenting theory, new algorithms, accurate retrospective designs, new prospective designs, and biochemical validation.

**Author summary:** Computational structure-based protein design is an innovative tool for redesigning proteins to introduce a particular or novel function. One such possible function is improving the binding of one protein to another, which can increase our understanding of biomedically important protein systems toward the improvement or development of novel therapeutics. Herein we introduce two novel, provable algorithms, fries and *EWAK*\*, for more efficient computational structure-based protein design as well as their application to the redesign of the c-Raf-RBD:KRas protein-protein interface. These new algorithms speed up computational structure-based protein design while maintaining accurate calculations, allowing for larger, previously infeasible protein designs. Using fries and *EWAK*\* within the osprey suite, we designed the tightest known binder of KRas, an “undruggable” cancer target. This new variant of a KRas-binding domain, c-Raf-RBD, should serve as an important tool to probe the protein-protein interface between KRas and its effectors as work continues toward an effective therapeutic targeting KRas.

## Introduction

Computational structure-based protein design (CSPD) is an innovative tool that enables the prediction of protein sequences with desired biochemical properties (such as improved binding affinity). osprey (Open Source Protein Redesign for You) [1] is an open-source, state-of-the-art software package used for CSPD and is available at http://www.cs.duke.edu/donaldlab/osprey.php for free. osprey’s algorithms focus on *provably* returning the optimal sequences and conformations for a given input model. In contrast, as argued in [2–7], stochastic, non-deterministic approaches [8–10] provide no guarantees on the quality of conformations, or sequences, and make determining sources of error in predicted designs very difficult.

When using osprey, the input model generally consists of a protein structure, a flexibility model (e.g., choice of sidechain or backbone flexibility, allowed mutable residues, etc.), and an all-atom pairwise-decomposable energy function that is used to evaluate conformations. osprey models amino acid sidechains using frequently observed rotational isomers or “rotamers” [11]. Additionally, osprey can also model continuous sidechain flexibility [12–15] along with discrete and continuous backbone flexibility [16–19], which allow for a more accurate approximation of protein behavior [13, 16, 20–23]. The output produced by CSPD generally consists of a set of candidate sequences and conformations. Many protein design methods have focused on computing a global minimum energy conformation (GMEC) [14, 18, 24–28]. However, a protein in solution exists not as a single, low-energy structure, but as a thermodynamic ensemble of conformations. Models that only consider the GMEC may incorrectly predict biophysical properties such as binding [12, 20–23, 29–31] because GMEC-based algorithms underestimate potentially significant entropic contributions. In contrast to GMEC-based approaches, the *K*\* algorithm [12, 29, 30] in osprey models thermodynamic ensembles to provably and efficiently approximate the *K*\* score. The *K*\* score is a ratio of the Boltzmann-weighted partition functions for a protein-ligand complex that estimates the association constant, *K*_a_ (further detailed in the Section entitled “Computational materials and methods”). *BBK*\* [32] is the most recent improvement on the traditional *K*\* algorithm that allows for multi-sequence design. Previous algorithms [12, 27, 29, 30,33–35] that design for binding affinity using ensembles are linear in the size of the sequence space *N*, where *N* is exponential in the number of simultaneously mutable residue positions. *BBK*\* is the first provable ensemble-based algorithm to run in time sublinear in *N*, making it possible not only to perform *K*\* designs over large sequence spaces, but also to enumerate a gap-free list of sequences in order of decreasing *K*\* score.

Osprey has been used successfully on several empirical, prospective designs including designing enzymes [12, 16, 22, 29, 36], resistance mutations [2, 37, 38], protein-protein interaction inhibitors [30, 39], epitope-specific antibody probes [40], and broadly-neutralizing antibodies [41, 42]. These successes have been validated experimentally *in vitro* and *in vivo* and are now being tested in several clinical trials [43–45]. However, while osprey has been successful in the past, as the size of protein design problems grows (e.g., when considering a large protein-protein interface), enumerating and minimizing the necessary number of conformations and sequences to satisfy the provable halting criteria in previous *K*\*-based algorithms [12, 29, 30] becomes prohibitive (despite recent algorithmic improvements [32]). The entire conformation space can be monumental in size and heavily populated with energetically unfavorable sequences and conformations. *EWAK*\*, an Energy Window Approximation to *K*\*, seeks to alleviate some of this difficulty by restricting the conformations included in each sequence’s thermodynamic ensemble. *EWAK*\* *guarantees* that each conformational ensemble contains *all* of the lowest energy conformations within an energy window of the GMEC for each design sequence. fries, a Fast Removal of Inadequately Energied Sequences, also mitigates this complexity problem by limiting the input sequence space to only the most favorable, low energy sequences. Previous algorithms have focused on optimizing for sequences whose conformations are similar in energy to that of the GMEC. In contrast, fries focuses on optimizing for sequences with energies better-than or comparable-to the wild-type sequence. fries *guarantees* that the restricted input sequence space includes all of the sequences within an energy window of the wild-type sequence, but excludes any potentially unstable sequences with significantly worse partition function values. Wild-type sequences are generally expected to be near-optimal for their corresponding folds [46]. Therefore, limiting the sequence space to sequences energetically similar to or better than the wild-type sequence is reasonable. A simplified workflow for fries/*EWAK*\* is presented in Fig 1. Compared to the previous state-of-the-art algorithm *BBK*\*, fries and *EWAK*\* improve runtimes by up to 2 orders of magnitude, fries decreases the size of the sequence space by up to more than 2 orders of magnitude, and *EWAK*\* decreases the number of conformations included in partition function calculations by up to almost 2 orders of magnitude.

**Fig 1.**
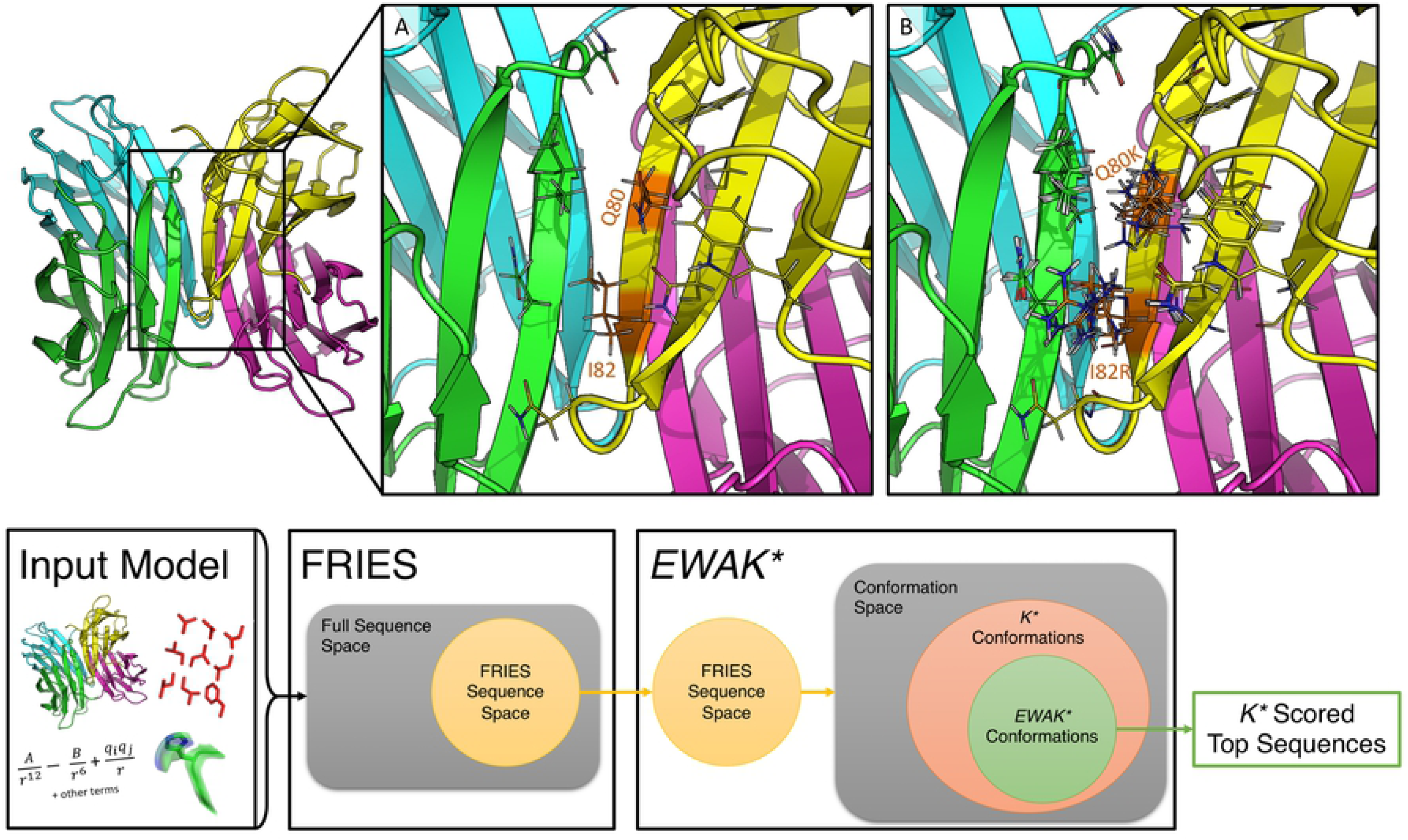
Design example using the structure of the LecB lectin *Pseudomonas aeruginosa* strain PA14 (PDB ID: 5A6Y [47]) and the OSPREY workflow for fries/*EWAK*\*. In the top panel, the full, 4 domain structure of lectin is shown on the left-hand side. (A) Zooming in on the region where domains A (green) and D (yellow) interact, showing the two mutable residues (Q80 and I82) along with the surrounding flexible shell of residues as lines. There were 11 flexible residues included in this design with Q80 and I82 allowed to mutate to all other amino acids except for proline. This design consisted of 8.102 ×10^11^ conformations and 441 sequences. fries limited this space to 5.704 ×10^11^ conformations and 206 sequences. fries/*EWAK*\* in combination reduced the amount of time taken by about 75% compared to *BBK*\*. fries alone was responsible for roughly 50% of this speed-up. (B) 10 low-energy conformations included in the thermodynamic ensemble of the design sequence with mutations Q80I and I82F. For this particular sequence, *BBK*\* minimized 10,664 conformations while *EWAK*\* minimized only 4,104 conformations. The bottom panel shows the general workflow for fries/*EWAK*\*. The workflow begins with the input model (as described in the Section entitled “Introduction”), which defines the design space for the first algorithm, fries. fries proceeds to prune the sequence space as described in the Section entitled “Fast Removal of Inadequately Energied Sequences (fries)” and as illustrated in the Venn diagram with the unpruned space shown as a yellow disk. Next, the remaining fries sequence space defines the conformation space (which contains multiple sequences as well as conformations) searched with *EWAK*\*. *EWAK*\* limits the conformations included in each partition function as described in the Section entitled “Energy Window Approximation to K* (*EWAK*\*).” *EWAK*\* generally searches over only a subset of the conformations (green area) that previous *K*\*-based algorithms like *BBK*\* [32] search (orange area). *EWAK*\* then returns the top sequences based on decreasing *K*\* score.

As a proof of concept to test these algorithms and our design approach, we used fries and *EWAK*\* to study the protein-protein interface (PPI) of KRas^GTP^ in complex with its tightest-binding effector, c-Raf. As described in the Section entitled “Computational redesign of the c-Raf-RBD:KRas protein-protein interface,” KRas is an important cancer target that has historically been considered “undruggable” [48]. Deepening the understanding of the PPI between KRas and its effectors is an important step toward developing effective new therapeutics. For this study, we focused on the redesign of the c-Raf Ras-binding domain (c-Raf-RBD) in complex with KRas^GTP^ (c-Raf-RBD:KRas^GTP^). First, our new algorithms successfully retrospectively predicted the effect on binding of mutations in the c-Raf-RBD:KRas^GTP^ PPI even where other computational methods previously failed [49]. Next, we used fries/*EWAK*\* prospectively to predict the effect of novel, previously unreported mutations in the PPI of the c-Raf-RBD:KRas^GTP^ complex. We then measured the binding of top osprey-predicted c-Raf-RBD variants to KRas using a bio-layer interferometry (BLI) assay with a single-concentration screen. This screen suggested that one of our new computationally-predicted c-Raf-RBD variants – c-Raf-RBD(Y), a c-Raf-RBD variant that includes the mutation V88Y – exhibits improved binding to KRas^GTP^. Next, we created a c-Raf-RBD variant, c-Raf-RBD(RKY), that included this new mutation, V88Y, together with two previously reported mutations [49], N71R and A85K. fries/*EWAK*\* computationally predicted that c-Raf-RBD(RKY) would bind more tightly to KRas^GTP^ than any other variant. The single-concentration screen using BLI also suggested that c-Raf-RBD(RKY) binds more tightly to KRas^GTP^ than the previously reported best variant [49]. The *K*_*d*_ values for the most promising variants were measured using a BLI assay with titration which confirmed our computational predictions and that, to the best of our knowledge, the novel construct c-Raf-RBD(RKY) is the highest affinity variant ever designed, with single-digit nanomolar affinity for KRas^GTP^.

## Computational materials and methods

The *K*\* algorithm’s [12, 29, 30] *K*\* score serves as an estimate of the binding constant, *K*_a_, and is calculated by first approximating the Boltzmann-weighted partition function of each state: unbound protein (*P*), unbound ligand (*L*), and the bound protein-ligand complex (*C*). Each Boltzmann-weighted partition function *Z*_*x*_ (**s**), *x* ∈ {*P, L, C*}, is defined as:

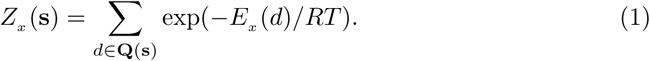

If **s** is any – generally amino acid – sequence of *n* residues, then **Q**(**s**) is the set of conformations defined by **s**, *E*_*x*_ (*d*) is the minimized energy of a conformation *d* in state *x*, and *R* and *T* are the gas constant and temperature, respectively. Many protein design algorithms approximate these partition functions for each state using either stochastic [50–53] or provable [2, 12, 29–31, 33, 53] methods.

Osprey’s *K*\* algorithm provably approximates these partition functions to within a user-specified *ε* of the full partition function as defined in Eq (1). The binding affinity for sequence **s** is defined as:

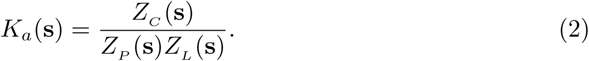

The *K*\* algorithm provably approximates this binding affinity. This is enabled by the use of *A*\* [4, 12, 26, 54], which allows for the gap-free enumeration of conformations in order of increasing lower bounds on energy [26]. However, enumerating a sufficient number of these conformations to obtain a guaranteed *ε*-approximation can be very time consuming because the set of all conformations **Q**(**s**) grows exponentially with the number of residues *n*. Also, the *K*\* algorithm was originally [12, 29, 30] limited to computing a *K*\* score for every sequence in the sequence space as defined by the input model for a particular design. However, *BBK*\* [32] builds on *K*\* and provably returns the top *m* sequences along with their *ε*-approximate *K*\* scores and runs in time sublinear in the number of sequences. That is, *BBK*\* does not require calculating *ε*-approximate *K*\* scores for (or even examining) every sequence in the sequence space before it returns the top sequences. Nevertheless, *BBK*\* may spend unnecessary time and resources evaluating unfavorable sequences before deciding to prune them.

To overcome the above limitations of *BBK*\* and *K*\*, we introduce fries, a Fast Removal of Inadequately Energied Sequences, and *EWAK*\*, an Energy Window Approximation to *K*\*. These two algorithms focus on limiting the input sequence space and the number of conformations included in each partition function estimate when approximating a sequence’s *K*\* score to provably only the most energetically favorable options. The fries/*EWAK*\* approach limits the number of conformations that must be enumerated (see the Section entitled “*EWAK*\* limits the number of minimized conformations when approximating partition functions while maintaining accurate K* scores”), which leads to significant speed-ups (see the Section entitled “fries/*EWAK*\* is up to 2 orders of magnitude faster than BBK*”) because each enumerated conformation must undergo an energy minimization step. This minimization step is relatively expensive, therefore, anything that reduces the number of minimized conformations while not sacrificing provable accuracy is desirable. For the importance of this minimization step to biological accuracy, see the discussions of continuous flexibility and its comparison to discrete flexibility in [4, 5, 7, 13, 14, 19]. *EWAK*\* also maintains the advances made by *BBK*\* including running in time sublinear in the number of sequences *N* and returning sequences in order of decreasing *K*\* score. fries and *EWAK*\* are described in further detail in the Section entitled “Algorithms” below.

### Algorithms

#### Fast Removal of Inadequately Energied Sequences (FRIES)

Generally in protein design when optimizing a protein-protein interface (PPI) for affinity, the designer aims to improve the *K*\* score of a variant sequence relative to the wild-type sequence, and, when performing a design targeting a similar fold, to minimally perturb the native structure. To accomplish this, fries guarantees to only keep sequences whose partition function values are not markedly worse than the wild-type sequence’s partition function values for all of the design states (e.g. protein, ligand, and complex). How many orders of magnitude worse a particular sequence’s partition function values are allowed to be is determined by a user-specified value *m*. The fries algorithm prunes sequences that exhibit massive decreases in partition function values that signal an increased risk of disturbing the native structure of the states in a given system. However, sequences with markedly worse, lower partition function values may be required when searching for, for example, resistance mutations, where positive and negative design are necessary [2, 37, 38]. Importantly, fries does still allow for sequences that may have lower, worse partition function values by allowing the user to specify how many orders of magnitude lower a candidate sequence’s partition function is allowed to be relative to the wild-type sequence’s partition function.

To prune the input sequence space, fries exploits *A*\* over a *multi-sequence tree* (as is described and used in comets [55]), which enjoys a fast sequence enumeration in order of lower bound on minimized energy. Each sequence *v* in this *multi-sequence tree* [55] has a corresponding *single-sequence conformation tree*, viz., a tree that can be searched for the lowest energy conformations for a sequence *v*. fries first enumerates sequences (in order of energy lower bounds) in the *multi-sequence tree* until the wild-type sequence is found. Then, fries searches the wild-type’s corresponding *single-sequence conformation tree* using *A*\*. The first conformation enumerated according to monotonic lower bound on pairwise minimized energy is then subjected to a full-atom minimization [30] to calculate the minimized energy of one of the wild-type sequence’s conformations *E*_*W T*_. fries then continues enumerating sequences in the *multi-sequence tree* in order of increasing lower bound on minimized energy until the lower-bound on the energy of a sequence *v*, 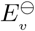, is greater than *E*_*W T*_ + *w* where *E*_*W T*_ is as described above and *w* is a user-specified energy window value (Fig 2). Any variant sequence *v* with a lower bound on minimized energy 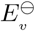 not satisfying the following criterion is pruned:

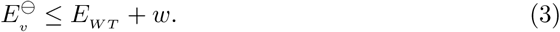

**Fig 2.**
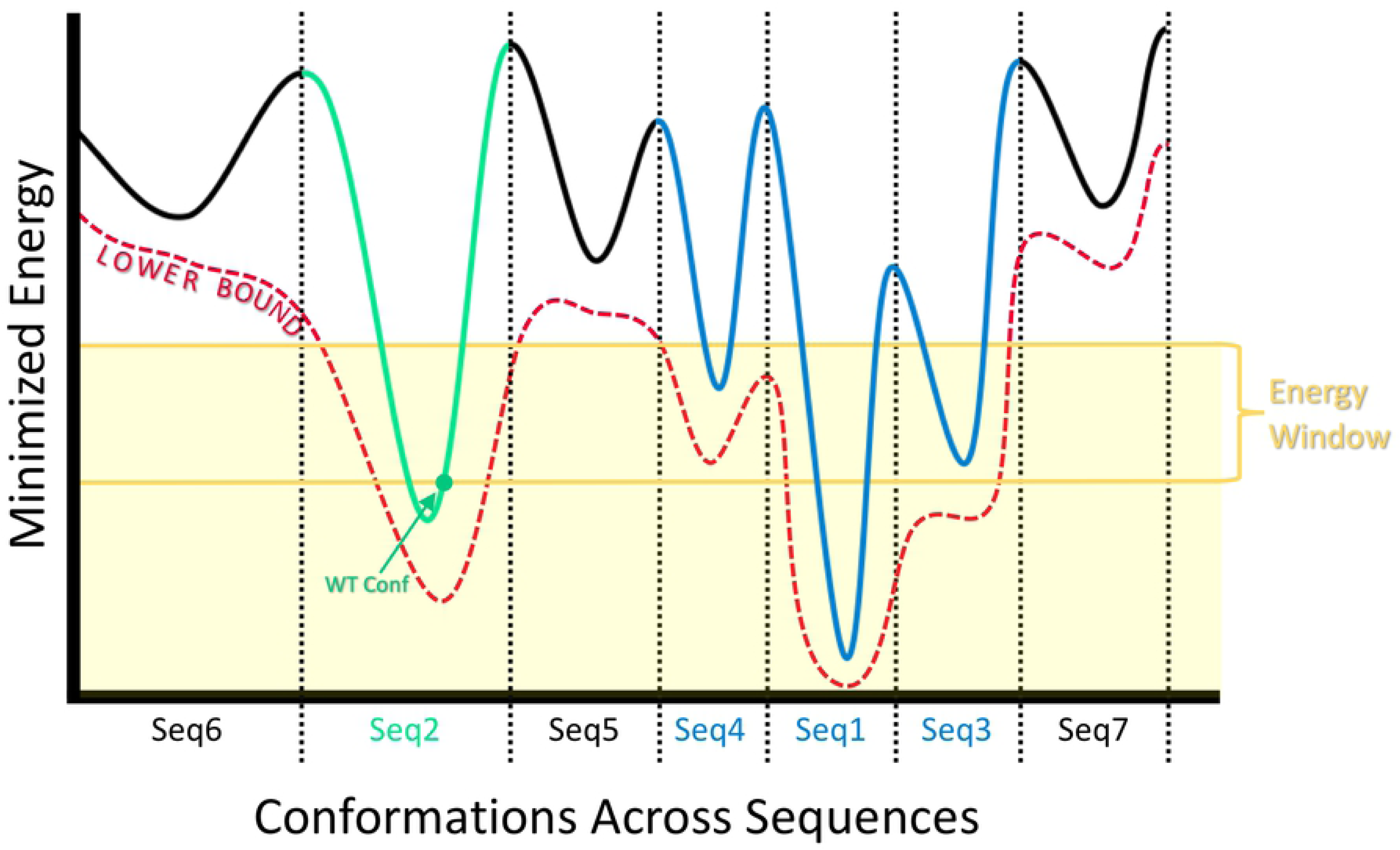
How fries chooses which sequences to keep and which sequences to prune. The solid curve represents the energy landscape of the conformation space that spans across, in this example, 7 different sequences (separated by dotted lines). Each sequence is labeled on the *x*-axis with an index indicating the order with which it is (or would be) enumerated with fries in order of increasing lower bound on minimized energy (red dotted curve). fries continues to enumerate in this way until it encounters the wild-type sequence (green), at which point fries calculates the minimized energy *E*_*W T*_ of the conformation with the lowest lower bound on minimized energy for the wild-type sequence (marked with a green dot). *E*_*W T*_ then becomes the baseline from which fries can provably enumerate all remaining sequences within some user-specified energy window *w* (yellow lines). Finally, fries prunes the sequences with energies provably higher than *E*_*W T*_ + *w* (black) and keeps the sequences that occur within the shaded yellow region (colored in blue and green). More sequences are also pruned according to their partition function values as described in the Section entitled “Fast Removal of Inadequately Energied Sequences (fries)” and as defined by Eq (4).

This criterion guarantees that the remaining, unpruned sequence space includes all sequences within an energy window of the wild-type sequence’s energy. fries enumerates sequences in order of increasing lower bound on minimized energy. Therefore, it calculates an upper bound 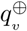 on the partition function for each sequence *v* by Boltzmann-weighting the lower bound on its energy 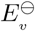 and multiplying it by the size of the conformation space for that particular sequence *|***Q**(*v*)*|*:

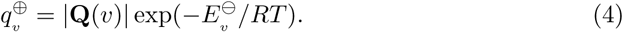

The lower bound for the wild-type sequence 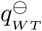 is calculated by Boltzmann-weighting the minimized energy of the single conformation found during the sequence search for the wild-type sequence *E*_*W T*_ :

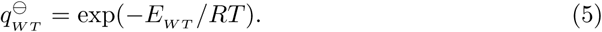

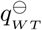 is a lower bound because, in the worst case, at least this one conformation will contribute to the partition function for the wild-type sequence. fries then uses these bounds to remove all of the sequences whose partition function value is not within some user-specified *m* orders of magnitude of the lower bound on the wild-type partition function 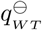. If the following criterion is not met, the sequence *v* is pruned from the space:

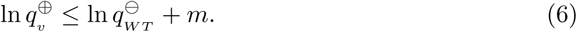

fries prunes sequences for the protein, the ligand, and the protein-ligand complex independently, limiting the input sequence space to exclude unfavorable sequences for all of the states. The resulting smaller sequence space is subsequently used as input for *EWAK*\*. The set of sequences remaining is guaranteed to include all of the sequences within a user-specified energy window *w* of the wild-type sequence that also satisfy the partition function criterion given in Eq (4). Importantly, fries can be used to limit the size of the input sequence space in this fashion for any of the protein design algorithms available within osprey.

#### Energy Window Approximation to K* (*EWAK*\*)

After reducing the size of the input sequence space using fries, as described in the Section entitled “Fast Removal of Inadequately Energied Sequences (fries),” *EWAK*\* proceeds by using a variation on an existing algorithm: *BBK*\* (described in [32]). The crucial difference between *BBK*\* and *EWAK*\* is that with *EWAK*\* the ensemble of conformations used to approximate each *K*\* score is limited to those within a user-specified energy window of the GMEC for each sequence. This guarantees to populate the partition function for a particular sequence and state with all of the provably lowest, most-favorable conformations (that fall within the user-specified energy window). These conformations often account for the majority of the full *ε*-approximate partition function (see the Section entitled “Computational materials and methods”) in traditional *K*\* calculations [12]. Hence, *EWAK*\* also empirically enjoys negligible loss in accuracy of *K*\* scores (see the Sections entitled “*EWAK*\* limits the number of minimized conformations when approximating partition functions while maintaining accurate K* scores” and “fries/*EWAK*\* retrospectively predicted the effect mutations in c-Raf-RBD have on binding to KRas”). *EWAK*\* retains the beneficial aspects of *BBK*\*, including returning sequences in order of decreasing predicted binding affinity and running in time sublinear in the number of sequences.

### Computational experiments

We implemented fries/*EWAK*\* in the osprey suite of open source protein design algorithms [1]. fries was tested on 2,662 designs that range from an input sequence space size of 441 to 10,164 total sequences. The size of the reduced input sequence space produced by fries was compared to the size of the full input sequence space size for each design. For these tests, fries returned every sequence within 8 kcal/mol of the wild-type sequence and was set to include only those sequences that are at most 2 orders of magnitude worse in partition function value than the wild-type. The results for these tests are described in the Section entitled “fries can reduce the size of the input sequence space by more than 2 orders of magnitude while retaining the most favorable sequences.” Computational experiments were also run comparing fries/*EWAK*\* with the previous state-of-the-art algorithm in osprey: *BBK*\* [32]. Using *BBK*\* and fries/*EWAK*\*, we computed the top 5 best binding sequences for 167 different designs to compare the running time of *BBK*\* vs. fries/*EWAK*\*. fries was limited to sequences within 4 kcal/mol of the wild-type sequence that are at most 2 orders of magnitude worse in partition function values than the wild-type. The *EWAK*\* partition function approximations were limited to conformations within an energy window of 1 kcal/mol of the GMEC for each sequence. *BBK*\* was set to return the top 5 sequences with an accuracy of *ε* = 0.68 (as was described in [32]). Using these same *EWAK*\* and *BBK*\* parameters, we also compared the change in the size of the conformation space necessary to compute an accurate *K*\* score for *BBK*\* vs. *EWAK*\* for 661 partition functions from 161 design examples. The results for these tests are described in Sections entitled “fries/*EWAK*\* is up to 2 orders of magnitude faster than BBK*” and “fries can reduce the size of the input sequence space by more than 2 orders of magnitude while retaining the most favorable sequences.” The number of conformations that undergo minimization (as described in [12–15]) for each partition function calculation with *EWAK*\* was also compared across different energy window sizes for 350 partition function calculations from 87 design examples. These partition function calculations were compared to *BBK*\*’s partition function calculations with a demanded accuracy of *ε* = 0.10. This smaller *ε* allowed for more accurate approximations of the *K*\* scores. The results for these tests are described in the Section entitled “fries can reduce the size of the input sequence space by more than 2 orders of magnitude while retaining the most favorable sequences.”

Every design included a set of mutable residues along with a set of surrounding flexible residues (Fig 1 for an example). All of these residues were allowed to be continuously flexible [12–15]. The designs were selected from 40 different protein structures (listed in S1 Table and also used in [32, 56]), and were run on 40-48 core Intel Xeon nodes with up to 200 GB of memory.

## Computational results

### fries can reduce the size of the input sequence space by more than 2 orders of magnitude while retaining the most favorable sequences

The number of remaining sequences after fries was compared to the size of the complete input sequence space. In the best case, when using fries, the sequence space was decreased by more than 2 orders of magnitude and the conformation space was decreased by just over 4 orders of magnitude. The sequence space was reduced an average of 49% and the conformation space was reduced an average of 40%. These results are broken down further in Fig 3.

**Fig 3.**
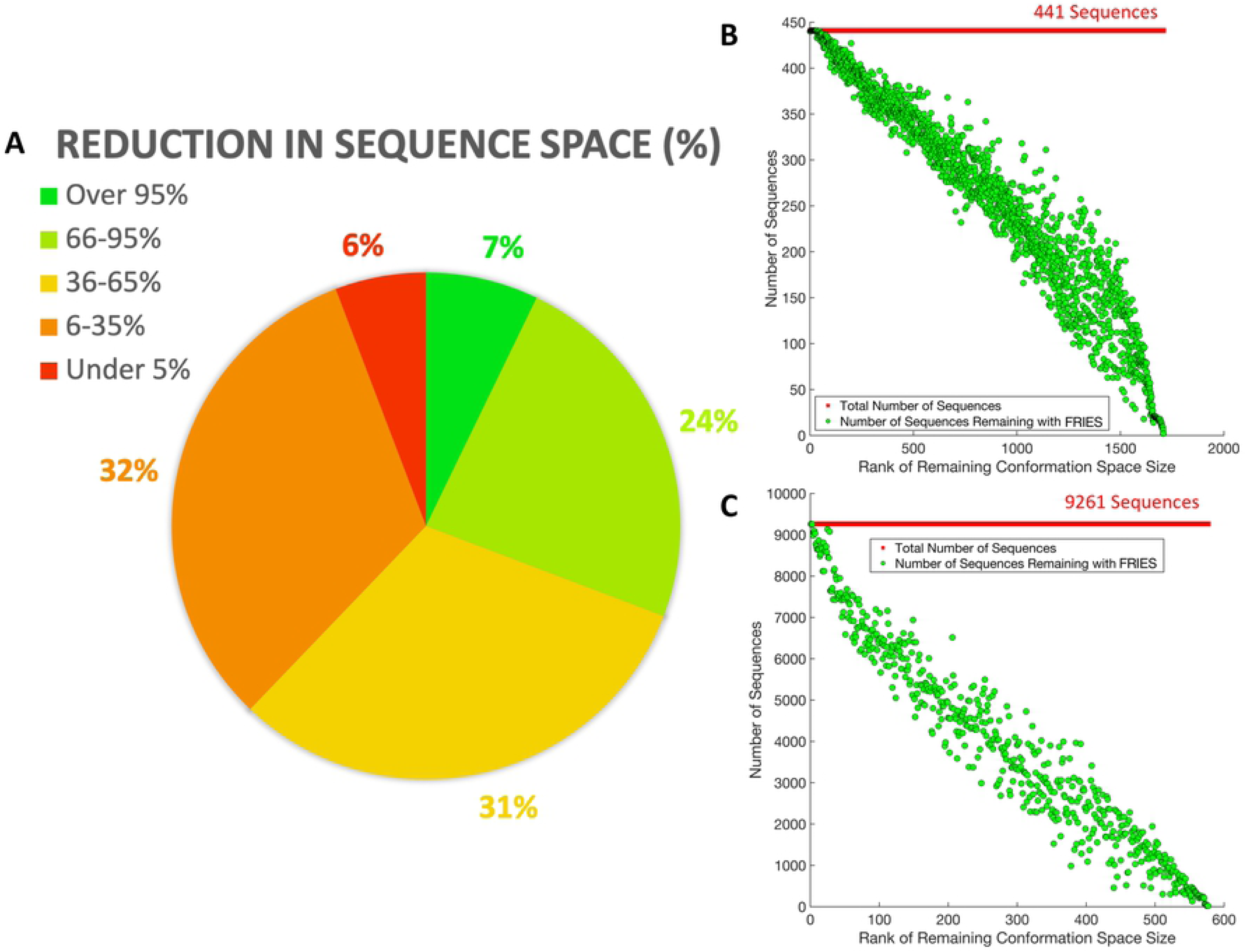
Reduction in input sequence space size using fries. (A) A pie chart representing the reduction in the sequence space in percentages across all 2,662 designs. 7% of the designs had a reduction in sequence space over 95%, 24% of the designs had a reduction in sequence space between 66-95%, 31% of the designs had a reduction in sequence space between 36-65%, 32% of the designs had a reduction in sequence space between 6-35%, and 6% of the designs had a reduction in sequence space under 5%. (B) and (C) plot the number of sequences remaining after using fries starting with 441 and 9,261 sequences total, respectively. The number of sequences remaining for each design are sorted in order of decreasing size of the remaining conformation space after fries.

### fries/*EWAK*\* is up to 2 orders of magnitude faster than BBK*

The overall runtime was compared between *BBK*\* and fries/*EWAK*\*. fries/*EWAK*\* was an average of 62% faster than *BBK*\* on 167 example design problems. fries removed unfavorable sequences (as described in the Section entitled “Fast Removal of Inadequately Energied Sequences (fries)”) from the search space for 156 out of the 167 design problems. For the cases described in the Section entitled “Computational experiments,” fries/*EWAK*\* performed consistently faster than *BBK*\* (in 92% of the design examples) as shown in Fig 4, Panel A. The longest running *BBK*\* design problem took nearly 8 days, whereas fries/*EWAK*\* completed the same example in just under 2 hours. In contrast, the design problem that took the longest for fries/*EWAK*\* out of the 167 tested only required about 22 hours (the same design took *BBK*\* over 178 hours).

**Fig 4.**
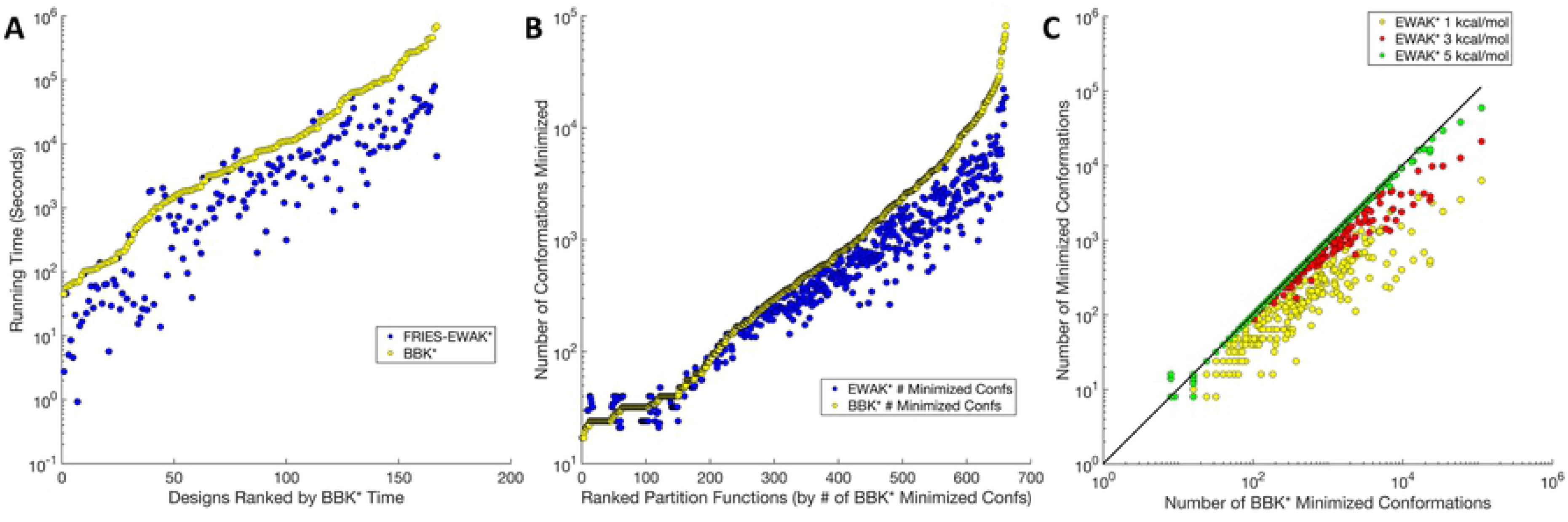
Comparing runtimes and the number of minimized conformations between fries/*EWAK*\* and *BBK*\* for a variety of designs. (A) A plot of the runtime in seconds (the *y*-axis is on a log scale) for fries/*EWAK*\* (blue dots) and *BBK*\* (yellow dots) for 167 design examples. Each point represents one design and is plotted in increasing order of *BBK*\* running time. fries/*EWAK*\* was faster than *BBK*\* 92% of the time with an average improvement of 62% over *BBK*\* and a maximum improvement of 2.2 orders of magnitude. This improvement was evident in (A) since the blue dots (fries/*EWAK*\* times) fall mostly below the yellow dots (*BBK*\* times). (B) A plot of the number of conformations minimized (*y*-axis is on a log scale) for 661 partition function calculations from 161 design examples. The number of conformations minimized by *EWAK*\* (blue dots) was less than the number of conformations minimized by *BBK*\* (yellow dots) in 68% of these cases, as is evidenced by the blue dots landing mostly below the yellow dots. In the best case, *EWAK*\* decreased the number of conformations by 1.1 orders of magnitude. The average percent reduction in the number of minimized conformations was 27%. (C) Each dot represents a calculated partition function. Yellow dots are partition functions limited to within a 1.0 kcal/mol window of the GMEC, red dots are partition functions limited to a 3.0 kcal/mol window of the GMEC, and green dots are partition functions limited to within a 5.0 kcal/mol window of the GMEC. These dots are plotted according to the number of minimized conformations required for each corresponding *BBK*\* partition function calculation. The solid black line represents the number of *BBK*\* minimized conformations, so dots that fall below the black line represent examples that required fewer minimized conformations than with *BBK*\*. As they approach the 5.0 kcal/mol window, the dots begin to converge with the *BBK*\* line. However, as the number of *BBK*\* minimized conformations rises beyond ~ 10^4^, even the green dots drop below the *BBK*\* line.

### *EWAK*\* limits the number of minimized conformations when approximating partition functions while maintaining accurate K* scores

We examined 661 *K*\* score calculations, and concluded that the total number of conformations minimized to approximate the *K*\* score was decreased by an average of 27%. In the best case the number of conformations minimized to approximate the *K*\* score was decreased by 93%. These results are plotted in Fig 4, Panel B. Even though the partition function approximations were limited to a smaller conformation space with *EWAK*\*, the *K*\* scores did not differ by more than 0.2 orders of magnitude between *EWAK*\* and *BBK*\* for these 661 example *K*\* score calculations.

A total of 350 of these 661 partition functions were subsequently re-estimated using *BBK*\* with a more accurate, stringent *ε* value of 0.1 and using *EWAK*\* with varied energy windows: 1.0 kcal/mol, 3.0 kcal/mol, and 5.0 kcal/mol. We examined the number of conformations minimized for each complex partition function calculation across the examples. When using 1.0 kcal/mol, *EWAK*\* minimized up to 1.7 orders of magnitude fewer conformations (Fig 4, Panel C for more details). Despite this decrease in the number of included conformations, *EWAK*\* reported accurate *K*\* scores. The largest difference in scores between *BBK*\* and *EWAK*\* was 0.3 orders of magnitude. The accuracy of *EWAK*\* is explored further in the Section entitled “fries/*EWAK*\* retrospectively predicted the effect mutations in c-Raf-RBD have on binding to KRas.”

### Computational redesign of the c-Raf-RBD:KRas protein-protein interface

We previously showed, by investigating 58 mutations across 4 protein systems, that osprey can accurately predict the effect of mutations on PPI binding [1]. Herein, we tested the biological accuracy of the new modules fries and *EWAK*\* after adding them to osprey by applying them to a particular system of significant biomedical and pharmacological importance: c-Raf-RBD in complex with KRas. The c-Raf Ras-binding domain (c-Raf-RBD) is a small self-folding domain that does not include the kinase signaling domains normally present in c-Raf. The c-Raf-RBD normally binds to KRas when KRas is GTP-bound (KRas^GTP^). A c-Raf-RBD variant that has high affinity for KRas^GTP^ could be an important first step toward discovering a tool that disrupts the KRas:effector interaction. Despite the recent successes with inhibitors targeting mutantKRas(G12C) by trapping it in the inactive GDP-bound state [57–62] and their recent move to clinical trials [63], these inhibitors are susceptible to resistance in the form of up-regulation of guanine nucleotide exchange factors (GEFs) and nucleotide exchange [60] which both push KRas to remain in its GTP-bound state. An inhibitor of the interaction between KRas^GTP^ and its effectors is hypothesized to have the advantage of not being susceptible to these mechanisms of resistance because it would directly interrupt KRas signaling. Hence, to further verify the accuracy and utility of fries/*EWAK*\*, we focused on this important PPI between KRas^GTP^ and one of its many effectors, c-Raf. First, in the Section entitled “fries/*EWAK*\* retrospectively predicted the effect mutations in c-Raf-RBD have on binding to KRas,” we retrospectively investigated previously reported mutations in the c-Raf-RBD [49, 64, 65] and how they affect the binding of c-Raf-RBD to KRas. This retrospective study lays the groundwork for the prospective study we present that investigates novel mutations. So, following the retrospective study, we computationally redesigned the PPI using fries/*EWAK*\* in search of new c-Raf-RBD variants with improved affinity for KRas^GTP^ (see the Section entitled “Prospective redesign of the c-Raf-RBD:KRas protein-protein interface toward improved binding” for details). To perform these computational designs, we first made a homology model of c-Raf-RBD bound to KRas^GTP^ (see S1 Text for details).

### fries/*EWAK*\* retrospectively predicted the effect mutations in c-Raf-RBD have on binding to KRas

Each previously reported c-Raf-RBD variant [49, 64, 65] was tested computationally using fries/*EWAK*\* by calculating a *K*\* score, a computational approximation of *K*_*a*_, for each variant along with its corresponding wild-type sequence. A percent change in binding was then calculated by comparing the variant’s *K*\* score to the corresponding wild-type sequence’s *K*\* score. The log_10_ of this value was then calculated and normalized to the wild-type by subtracting 2. A similar procedure was completed using the reported experimental data in order to easily compare the computationally predicted effect with the experimentally measured effect. The resulting value, called Δb, represents the change in binding. If a variant has a Δb less than 0, it is predicted to decrease binding. If a variant has a Δb greater than 0, it is predicted to increase binding. Δb values that are roughly equivalent to 0 indicate variants that have little to no effect on binding since the wild-type sequence was normalized to 0. The Δb values for the 41 computationally tested variants were plotted and compared to experimental values in Fig 5.

**Fig 5.**
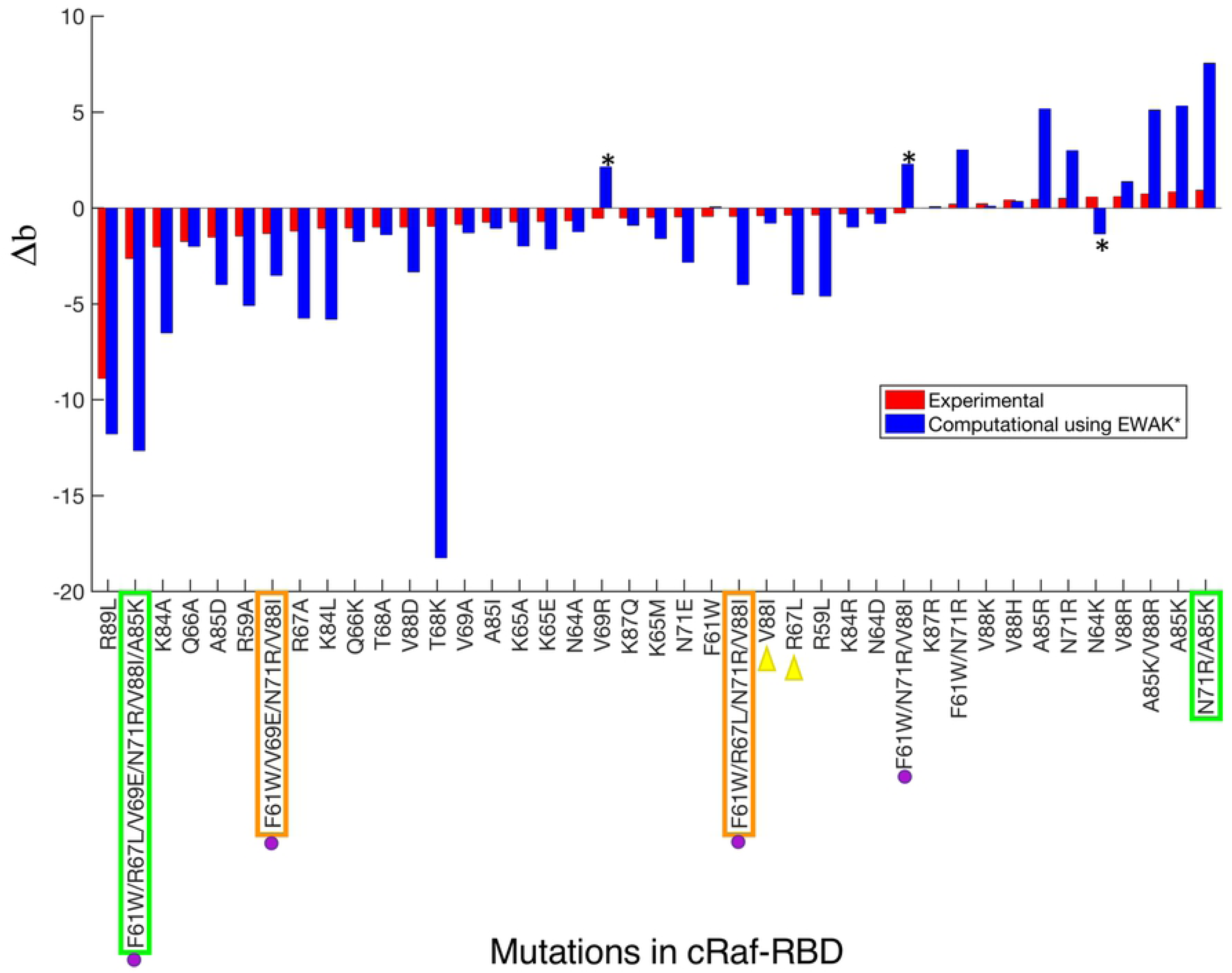
Predicting the effect of mutations in c-Raf-RBD on binding with KRas. Each bar represents either the experimental (red) or computationally predicted (blue) effect each variant has on binding. The bars are sorted in increasing order of Δb value (see the Section entitled “fries/*EWAK*\* retrospectively predicted the effect mutations in c-Raf-RBD have on binding to KRas”) of the experimental (red) bars. If the Δb value is less than 0, binding decreases. If the Δb value is greater than 0, binding increases. If the Δb value is close to 0, the effect is neutral. Quantitative values of *K*\* tend to overestimate the biological effects of mutations (leading to the much larger blue bars) due to the limited nature of the input model compared to a biologically accurate representation. However, *K*\* in general does a good job ranking variants, as can be seen here in Fig 6, in [1], and in [38]. Out of the 41 variants listed on the *x*-axis, only 3 were predicted incorrectly (marked with black asterisks) by *EWAK*\*. In terms of accuracy, *BBK*\* performed very similarly to *EWAK*\* (data not shown), however, in 2 cases (marked with green boxes), *BBK*\* ran out of memory and was unable to calculate a score. *BBK*\* also did not return values for the 2 variants marked with orange boxes. The variants marked with purple dots were tested in [49] experimentally – not computationally – and decreased binding of c-Raf-RBD to KRas^GTP^ was observed, which *EWAK*\* was able to predict correctly. The two variants marked with yellow triangles were computationally predicted in [49] to improve binding of c-Raf-RBD to KRas^GTP^. However, the experimental validation in [49] showed that these variants exhibit decreased binding, which *EWAK*\* accurately predicted.

Out of the 41 variants tested (see S2 Table), *EWAK*\* predicted the experimentally-reported effect (increased vs. decreased binding) correctly in 38 cases. The three designs where the effect was predicted incorrectly are marked with a star in Fig 5. To make these predictions, the corresponding computational designs ranged in size from single point mutations up to 6 simultaneous mutations. Results are outlined in Fig 5. Furthermore, the Spearman’s *ρ* value – a measure of the correlation between two sets of rankings – when comparing the experimental data to the computational predictions is 0.81. This *ρ* value indicates that not only can *EWAK*\* correctly predict the effect of a particular set of mutations, but that *EWAK*\* also does a good job ranking the variants in order according to change in binding upon mutation (Fig 6). This value is very similar to Spearman’s *ρ* values for other PPI systems when using osprey [1].

**Fig 6.**
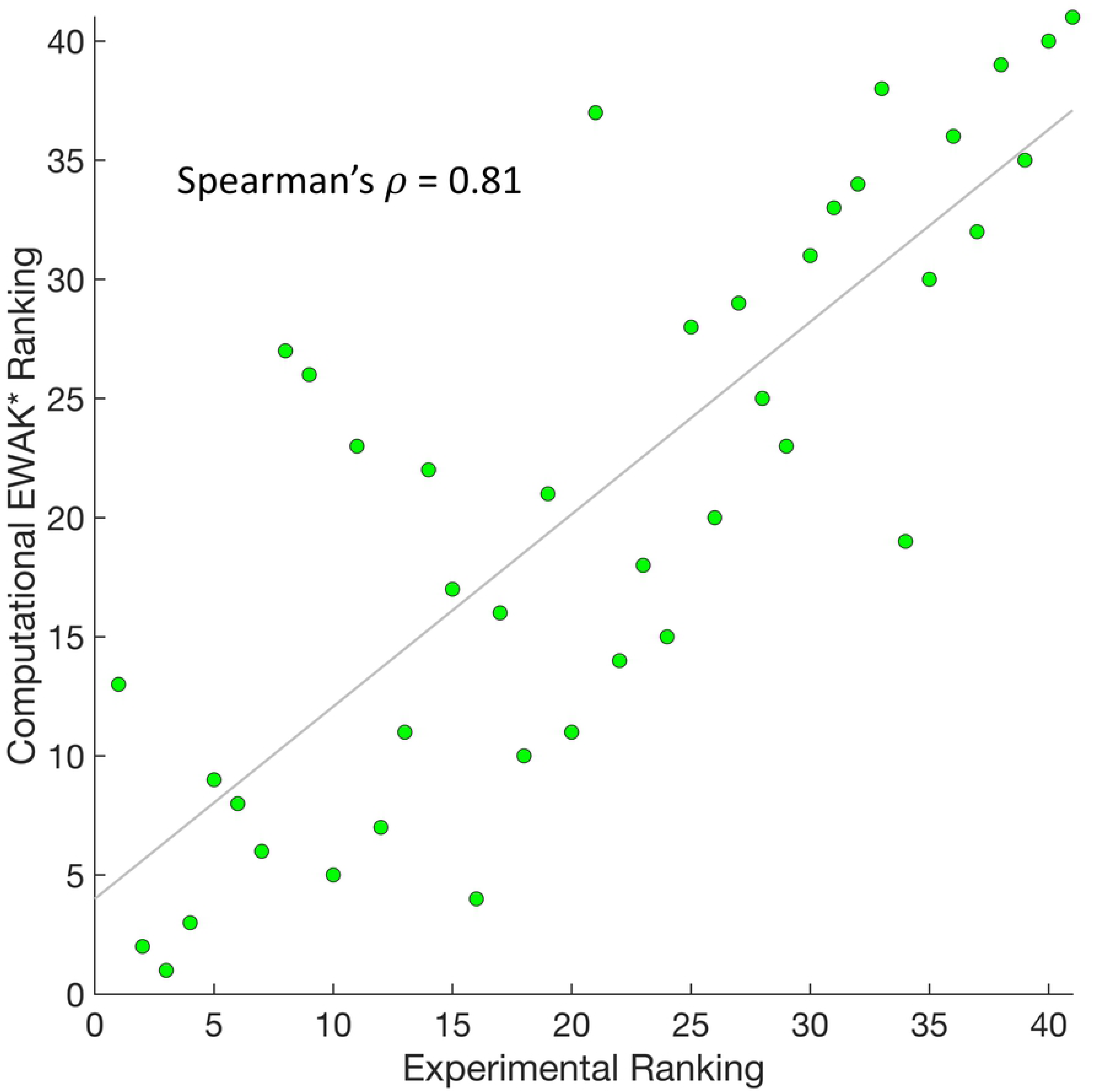
Comparing the computational *EWAK*\* ranking with the experimental ranking for 41 c-Raf-RBD variants binding to KRas. Each green dot represents a variant of c-Raf-RBD and is plotted according to the experimental ranking along with the corresponding computational ranking of its binding to KRas. A least squares fit line is shown in gray. Calculating the Pearson correlation coefficient between the two sets of rankings yields a Spearman’s *ρ* of 0.81.

*BBK*\* produced similarly accurate results, but took up to 10 times longer and failed to produce results in 4 cases. In particular, in 2 cases (marked in green in Fig 5), *BBK*\* ran out of memory. These cases serve as examples of large designs where *EWAK*\* outperforms *BBK*\*. In the 2 other cases (marked in orange in Fig 5), *BBK*\* failed to return a result for the requested sequence in the top 5 reported sequences. This illustrates how *EWAK*\* and fries are particularly helpful when performing larger designs that contain more simultaneous mutations and more flexible residues.

Finally, we compared our predictions to the interesting biological predictions in [49]. It is unclear how many mutants were computationally evaluated, but the authors do report computational predictions for 6 point mutations. Of those, point mutants R67L, N71R, and V88I were predicted to improve the intermolecular interactions between c-Raf-RBD and KRas^GTP^. However, experiments found that R67L and V88I actually reduced the binding of c-Raf-RBD to KRas^GTP^ [49, 64]. In contrast to [49], *EWAK*\* accurately predicted that these mutations decrease binding of c-Raf-RBD to KRas^GTP^. For a more detailed view of one of these designs, V88I, see Fig 7. Additionally, a number of mutations were combined and experimentally tested in [49]. Unfortunately, none of these variants improved binding to either KRas^GTP^ or KRas^GDP^, which fries/*EWAK*\* correctly predicted computationally (Fig 5). In [49], the authors do not present any computational predictions for these combined variants, but our results show that a computational prediction using osprey’s *EWAK*\* would have saved the time and resources taken to experimentally test these variants.

**Fig 7.**
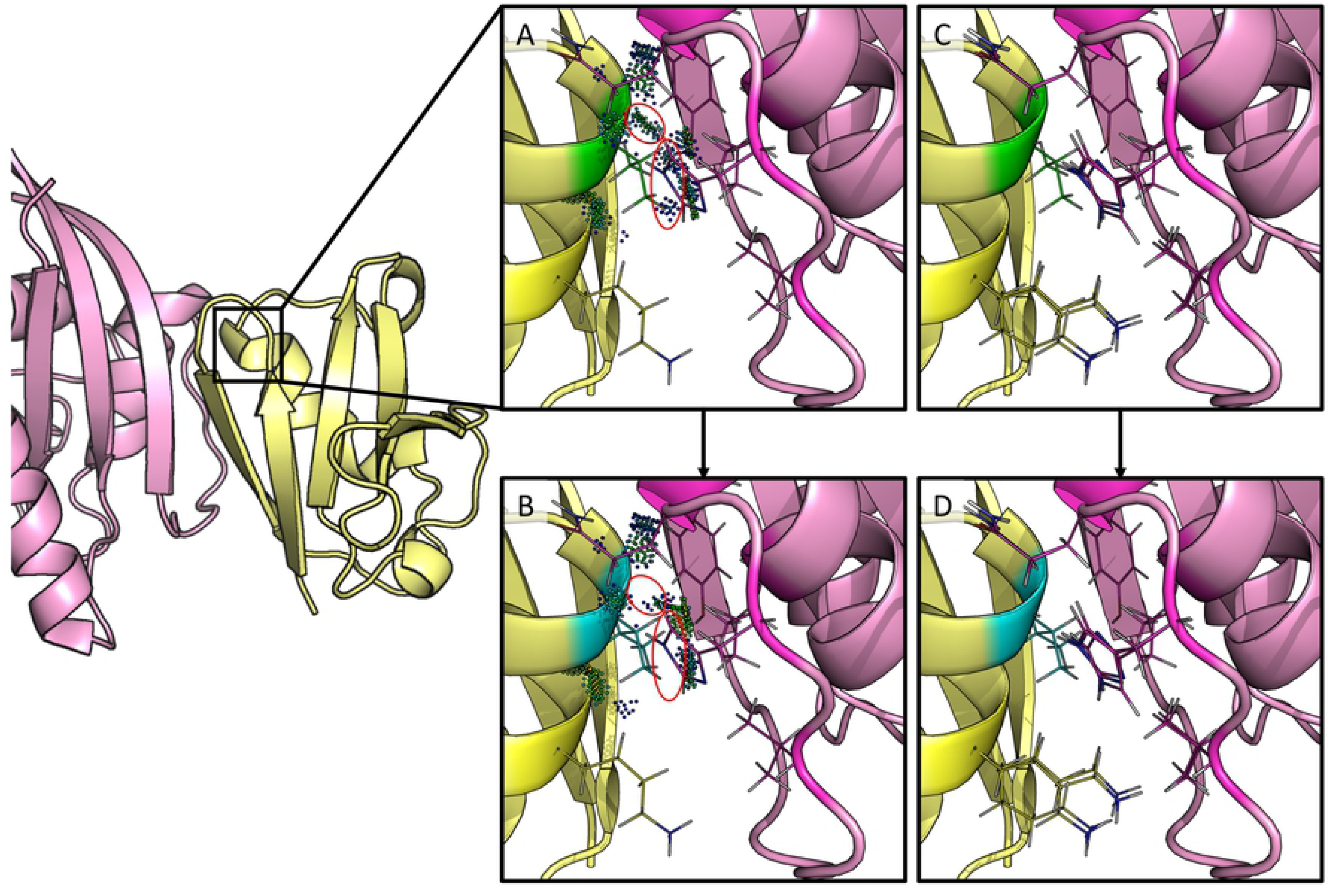
Redesign of c-Raf-RBD residue position 88 from valine to isoleucine. The left-hand side shows c-Raf-RBD (yellow) in complex with KRas (pink). Panels (A-D) zoom in on one particular design at residue position 88 and are rotated 180°. Residue position 88 has a valine in the native, wild-type sequence (panels A & C) which was redesigned to an isoleucine (panels B & D). A mutation to isoleucine at this position was computationally predicted by *EWAK*\* to decrease the binding of c-Raf-RBD to KRas^GTP^. This was experimentally validated in [49], where the authors incorrectly computationally predicted the effect of this particular mutation on the binding of c-Raf-RBD to KRas^GTP^. (A) The wild-type residue (valine) is shown in green with dots that indicate molecular interactions [66] with the surrounding residues (residues allowed to be flexible in the design are shown as lines). (B) The mutant residue (isoleucine) is shown in blue with dots that indicate molecular interactions [66] with the surrounding residues (residues allowed to be flexible in the design are shown as lines). Contacts made by the wild-type valine residue (circled dots in (A)) were lost upon mutation to isoleucine (circled space in (B)). (C & D) A set of 10 low-energy conformations that were included in the corresponding partition function calculation are shown for the wild-type (green) and the variant (blue).

### Prospective redesign of the c-Raf-RBD:KRas protein-protein interface toward improved binding

The ability to accurately predict the effect mutations have on the binding of c-Raf-RBD to KRas^GTP^ (see the Section entitled “fries/*EWAK*\* retrospectively predicted the effect mutations in c-Raf-RBD have on binding to KRas”) gave us confidence in the *EWAK*\* algorithm’s ability to predict new mutations in this interface toward a c-Raf-RBD variant that exhibits an even higher affinity for KRas^GTP^ than previously reported variants which focused on targeting KRas^GDP^ [49]. Therefore, to do a prospective study, we computationally redesigned 14 positions in c-Raf-RBD in the c-Raf-RBD:KRas PPI to identify promising mutations. After extending osprey to include fries and *EWAK*\*, 14 different designs were completed where each design included 1 mutable position that was allowed to mutate to all amino acid types except for proline. Each design also included a set of surrounding flexible residues within roughly 4 Å of the mutable residue. These designs were run using fries and *EWAK*\* and included continuous flexibility [12–15]. fries was first used to limit each design to only the most favorable sequences (as described in the Section entitled “Fast Removal of Inadequately Energied Sequences (fries)”) and then *EWAK*\* was used to estimate the *K*\* scores (as described in the Section entitled “Energy Window Approximation to K* (*EWAK*\*)”). We report the upper and lower bounds on the *EWAK*\* score for each design in Table 1 (also see S3 Table), where the listed sequences are those that were not pruned during the fries step. From these results, the predicted binding effect (increased vs. decreased) was determined based on comparing each variant’s *K*\* score to its corresponding wild-type *K*\* score. We then selected 5 novel point mutations – that to our knowledge are not reported in any existing literature – for experimental validation (Table 1). It is worth noting that these 5 point mutations were selected out of an initial 294 possible mutations. We limited our experimental validation to only these 5 new mutations and 2 previously reported mutations. This greatly reduced the amount of resources necessary for experimental validation compared to testing all 294 possibilities. These mutations were selected based on having a promising *K*\* score and through examining structures calculated by *EWAK*\*. Of the mutations selected, T57M was selected to act as a variant that was computationally predicted to be comparable to wild-type. This variant was included to further verify the accuracy of osprey’s predictions. On the other hand, some of osprey’s top predictions were excluded, for instance, T57R (included in S3 Table) was not selected for experimental testing because it has an unsatisfied hydrogen bond as evidenced in the structures calculated by osprey. Therefore, we do not believe that the score accurately represents the effect the mutation will have. Other excluded top predictions (see S3 Table) displayed similar characteristics or have been reported and tested previously [49, 64, 65].

**Table 1.**
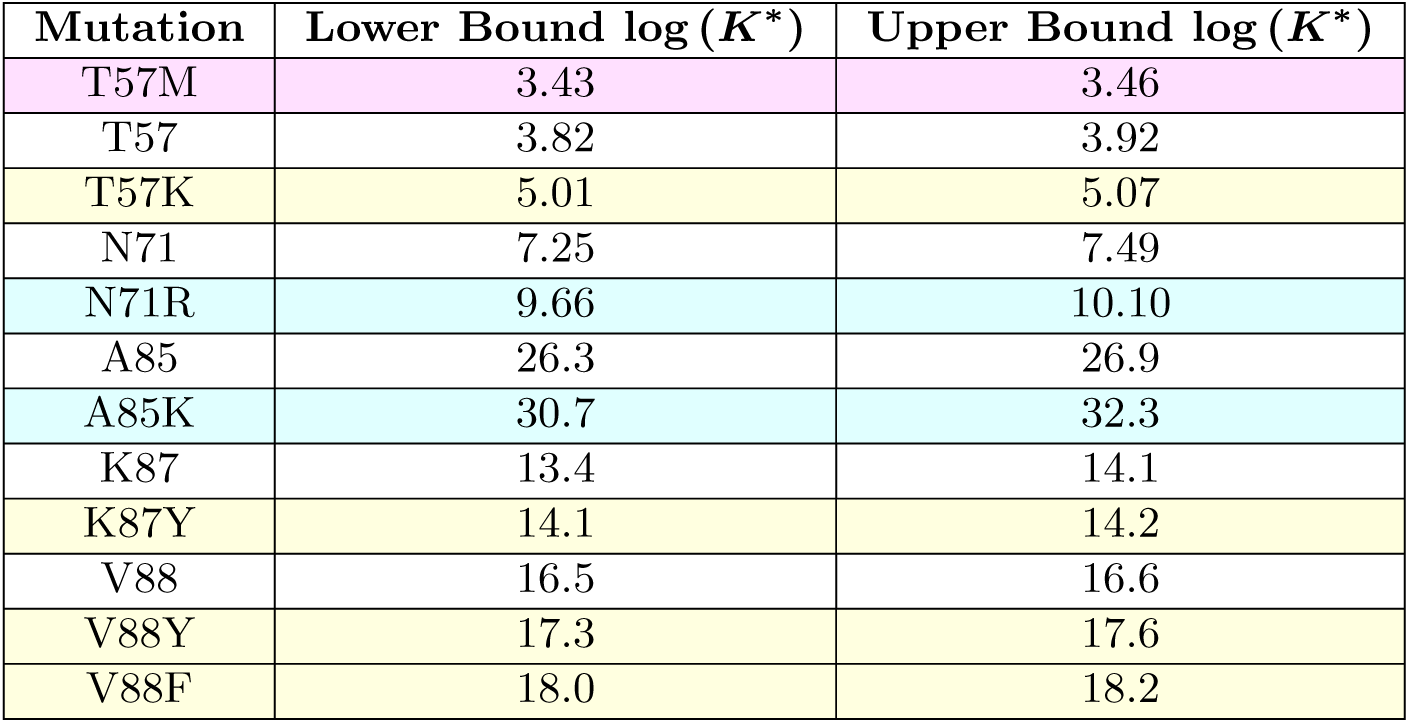
Computational predictions by OSPREY/fries/*EWAK*\* that were selected for experimental validation. Each row of the table shows the results of the redesign of a residue position in c-Raf-RBD in the c-Raf-RBD:KRas PPI that were also selected for experimental validation (all of the computational results are listed in S3 Table). The table contains the values for upper and lower bounds on log(*K*\*) values (the calculation of these bounds is described in detail in [32]). Mutations highlighted in yellow, blue, and pink were selected for experimental testing and validation. The two residues highlighted in blue are the best previously discovered [49] mutations that improve binding (independently and additively) and are included in our tightest binding variant, c-Raf-RBD(RKY) (Figs 9, 8, and 10). The variants highlighted in yellow are, to the best of our knowledge, never-before-tested variants that are predicted to increase the binding of c-Raf-RBD to KRas^GTP^. The variant highlighted in pink was selected for experimental testing to act as a mutation predicted to be comparable to wild-type to test how accurately osprey predicted the effects of these mutations.

| Mutation | Lower Bound $\log(K^*)$ | Upper Bound $\log(K^*)$ |
| --- | --- | --- |
| T57M | 3.43 | 3.46 |
| T57 | 3.82 | 3.92 |
| T57K | 5.01 | 5.07 |
| N71 | 7.25 | 7.49 |
| N71R | 9.66 | 10.10 |
| A85 | 26.3 | 26.9 |
| A85K | 30.7 | 32.3 |
| K87 | 13.4 | 14.1 |
| K87Y | 14.1 | 14.2 |
| V88 | 16.5 | 16.6 |
| V88Y | 17.3 | 17.6 |
| V88F | 18.0 | 18.2 |

### Experimental validation of mutations in the c-Raf-RBD:KRas protein-protein interface

The mutations selected (highlighted in Table 1) from computational design were experimentally validated using a bio-layer interferometry (BLI) assay. Results from an initial single-concentration BLI screen (Fig 8) suggested that, contrary to the computational predictions, the T57K and V88F variants decrease binding, whereas the T57M and K87Y mutations both have a roughly neutral effect on binding, which is consistent with the computational predictions. The final computationally predicted point mutant, V88Y, improves binding a comparable amount to the improvement seen with A85K or N71R, two previously reported variants also predicted by osprey and experimentally tested herein that improve binding. With the discovery of this new variant containing the point mutant V88Y (referred to herein as c-Raf-RBD(Y)) the next natural step was to combine it with the mutations found in the best reported variant, N71R and A85K (referred to herein as c-Raf-RBD(RK)). Therefore, we also included the double-mutant, c-Raf-RBD(RK), and the new triple-mutant – which contains N71R, A85K, and V88Y and is referred to herein as c-Raf-RBD(RKY) – in our initial BLI screen. Additionally, the c-Raf-RBD(RKY) variant was computationally predicted by fries/*EWAK*\* to bind to KRas^GTP^ more tightly than the previous best known binder, c-Raf-RBD(RK) (results are detailed in Fig 9). Given the promising screening and computational results for the c-Raf-RBD(Y) and c-Raf-RBD(RKY) variants, we measured *K*_*d*_ values for each variant by titrating the analyte over the ligand in a BLI-based assay (Fig 10). Excitingly, c-Raf-RBD(RKY) is calculated by the data from the BLI assay (Figs 8 and 10) to bind KRas^GTP^ roughly 5 times better than the previous best known binder, c-Raf-RBD(RK), and approximately 36 times better than wild-type c-Raf-RBD. Given how heavily studied the KRas system is, with several reported mutational and structural studies [49, 64, 64, 65, 65, 67, 67–73, 73, 74, 74–79], this is a discovery of some significance.

**Fig 8.**
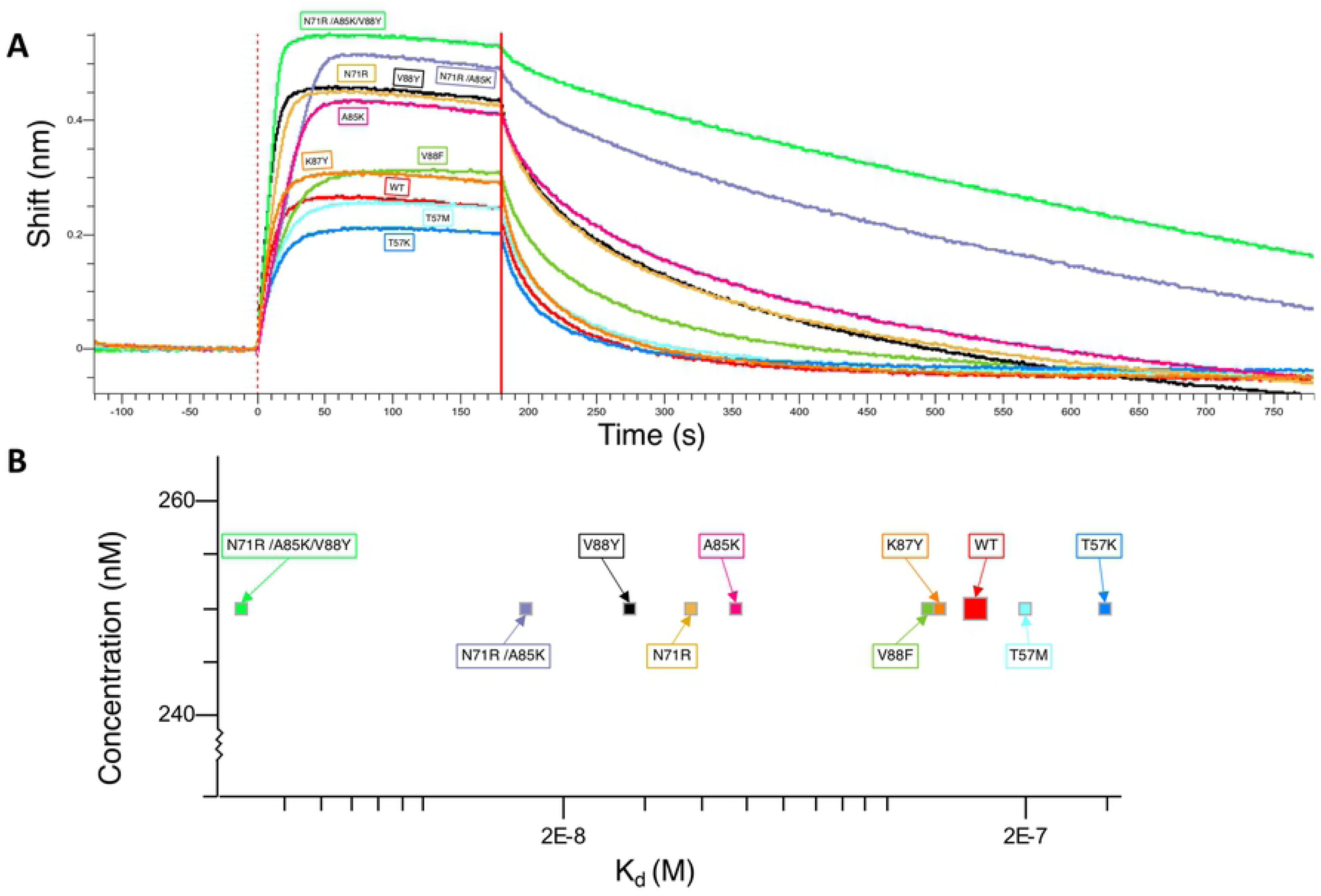
Single-concentration experimental screening of c-Raf-RBD variants binding to KRas using BLI. (A) Binding curves are shown for each variant (labeled on the plot) tested at a concentration of 250 nM. The colors and labels in panel (A) correspond to those in panel (B). (B) Plot of estimated *K*_*d*_ values for each tested variant from a single-concentration screen (plotted in panel (A)). The c-Raf-RBD(RKY) variant (in green on the far left) is a novel, newly discovered variant of c-Raf-RBD. Top variants were further validated and had their *K*_*d*_ values calculated more accurately using BLI titration experiments (Fig 10).

**Fig 9.**
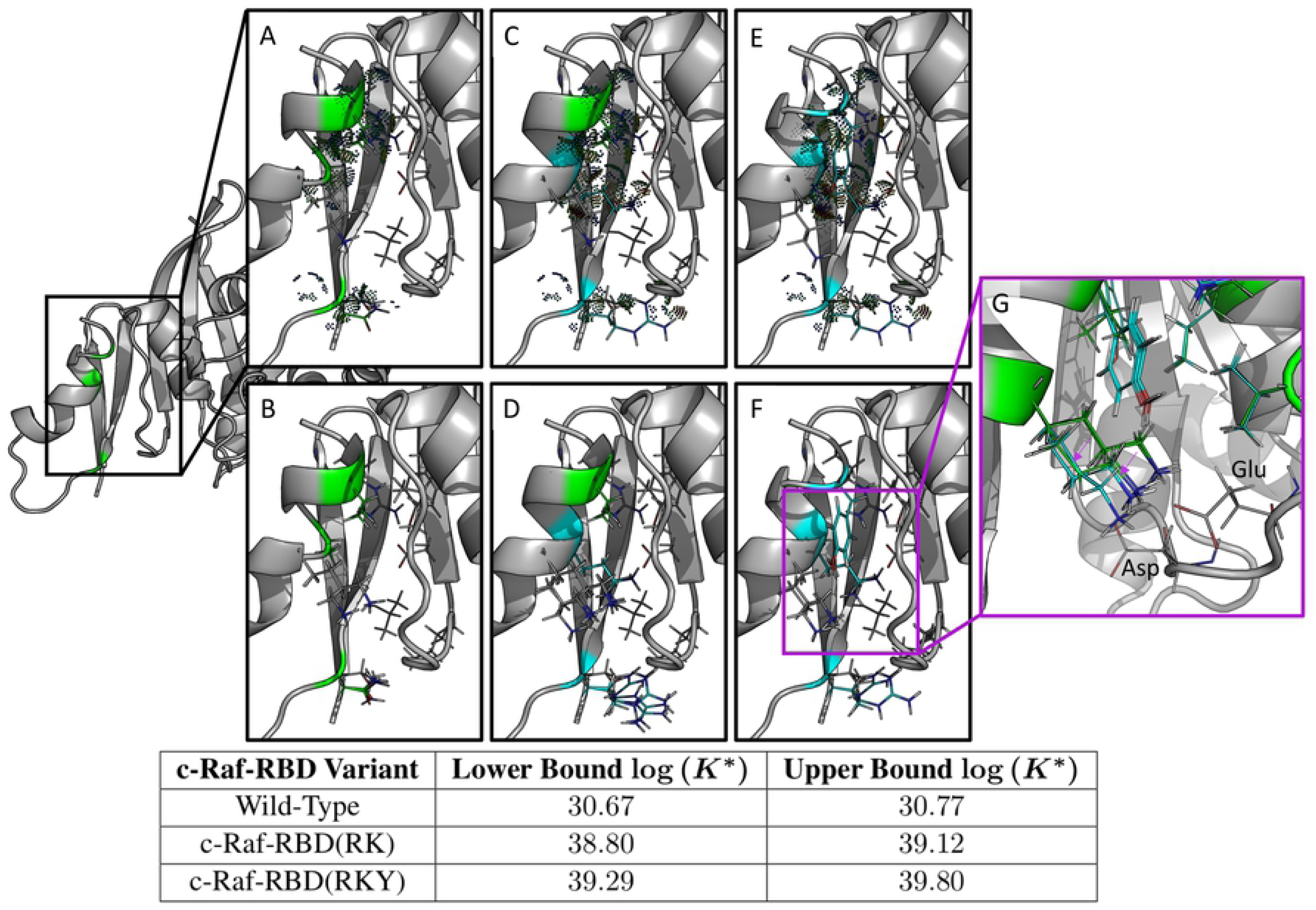
Computational predictions in the protein-protein interface of the c-Raf-RBD:KRas complex for c-Raf-RBD(RK) and the novel variant c-Raf-RBD(RKY). Shown on the left is only the relevant protein-protein interface between c-Raf-RBD and KRas. Each panel zooms in on this interface and details a different c-Raf-RBD variant and its corresponding computational predictions. The upper and lower bounds on the log(*K*\*) score for each design variant (wild-type, c-Raf-RBD(RK), and c-Raf-RBD(RKY)) are given in the bottom table. These computational predictions correspond with and are supported by the experimental results presented in the Section entitled “Experimental validation of mutations in the c-Raf-RBD:KRas protein-protein interface.” Panels (A) and (B) show the wild-type sequence, panels (C) and (D) show the variant c-Raf-RBD(RK), and panels (E) and (F) show the novel computationally predicted variant c-Raf-RBD(RKY). Panels (A), (C), and (E) show the wild-type, c-Raf-RBD(RK), and c-Raf-RBD(RKY), respectively, along with probe dots [66] that represent the molecular interactions within each structure calculated by osprey. These probe dots were selected to only show interactions between the residues included in the computational designs (shown as green and blue lines) with their surrounding residues. Panels (B), (D), and (F) show 10 low-energy structures from each conformational ensemble calculated by osprey/*EWAK*\*. Panel (G) shows a zoomed-in overlay of the wild-type variant with the c-Raf-RBD variant that includes only the V88Y mutation. Purple arrows indicate the change in positioning of the lysine at residue position 84 upon mutation of residue position 88 from valine to tyrosine. When valine is present at position 88, the lysine residue (shown in green) primarily hydrogen bonds with an aspartate (labeled) in KRas. When valine is mutated to tyrosine (shown in cyan), the lysine at position 84 moves to make room for the tyrosine and positions itself to hydrogen bond with both the aspartate and the glutamate (labeled) in KRas.

**Fig 10.**
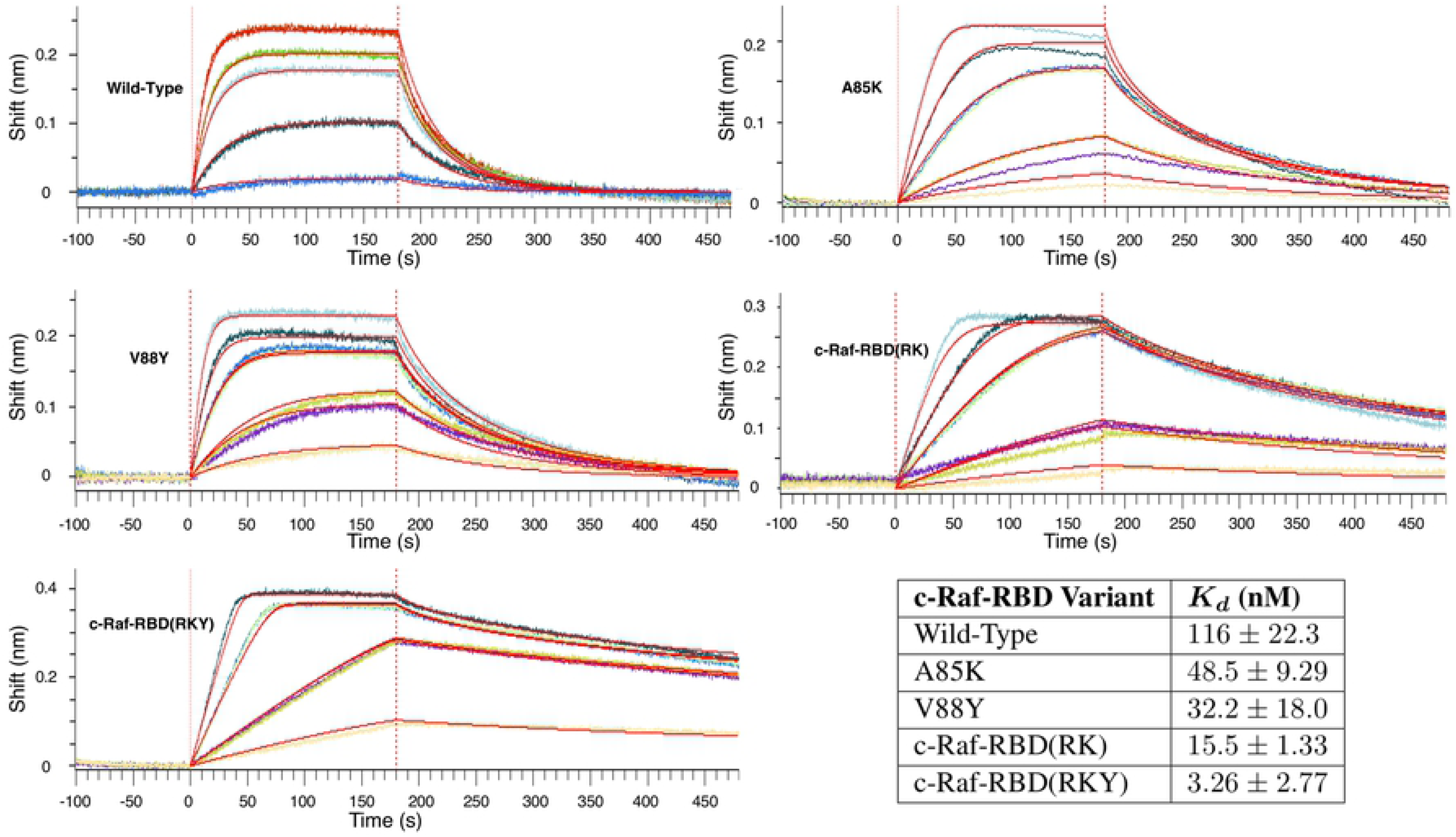
BLI titration experiments to calculate *K*_*d*_ values for select c-Raf-RBD variants. The plots shown here are representative and the data from replicate experiments are presented in S4 Table. Each plot shows the data collected from a titration BLI experiment where the concentration of the c-Raf-RBD variant is incrementally increased. The concentrations for the wild-type variant were 10, 50, 150, 200, and 300 nM. The concentrations for all of the other variants were 10, 25, 25, 75, 75, 125, and 200 nM. Repeat intermediate concentrations were used as loading controls. These curves were then fit using a mass transport model within the Octet Data Analysis HT software provided by FortéBio in order to calculate the *K*_*d*_ value for each variant’s binding to KRas. The values in the table here (bottom right) are average *K*_*d*_ values shown with 2 standard deviations calculated from replicate experiments (S4 Table). The values presented here for Wild-Type, A85K, and c-Raf-RBD(RK) agree well with previously reported *K*_*d*_ values [49]. The best binding variant, c-Raf-RBD(RKY), binds to KRas about 5 times better than the previous tightest-known binder, c-Raf-RBD(RK), and about 36 times better than wild-type c-Raf-RBD.

### Bio-layer interferometry (BLI) assay

Binding of wild-type and variants of c-Raf-RBD were experimentally measured using a bio-layer interferometry (BLI) assay. Each variant of c-Raf-RBD was expressed and purified (S2 Text) with cysteine residues at positions 81 and 96 substituted for isoleucine and methionine, respectively. These mutations were previously reported to minimally affect on the stability of c-Raf-RBD [73] and their substitution allows for the use of the c-Raf-RBD constructs in other assays (not mentioned herein). Additionally, we do not believe these residue substitutions have a large effect since the *K*_*d*_ values determined herein align with previously reported *K*_*d*_ values [49] (Fig 10). KRas was expressed and purified (S3 Text) with a poly-histidine protein tag (His-tag) and loaded with a non-hydrolyzable GTP analog, GppNHp. KRas was also made to include a substitution at position 118 from cysteine to serine in order to increase expression and stability [80]. Ni-NTA tips were then used to perform the BLI experiments to determine binding of the c-Raf-RBD variants to KRas^GppNHp^ (results are shown in Figs 8 and 10 and S4 Table). All experiments were carried out in 30 mM phosphate pH 7.4, 327 mM NaCl, 2.7 mM KCl, 5 mM MgCl_2_, 1.5 mM TCEP, 0.1% BSA, and 0.02% Tween-20 + Kathon at 25°C with 1000 RPM shaking and a KRas loading concentration of 20 µg/ml. Each curve presented (Figs 8 and 10) was fit using the built-in mass transport model within the Octet Data Analysis HT software provided by FortéBio. We only accepted fits with a sum of square deviations *χ*^2^ less than 1 (FortéBio recommends a value less than 3) and a coefficient of determination *R*^2^ greater than 0.98.

## Discussion

Fries and *EWAK*\* are new, provable algorithms for more efficient ensemble-based computational protein design. Efficiency and efficacy were tested and shown across a total of 2,826 different design problems. An implementation of fries/*EWAK*\* is available in the open-source protein design software OSPREY [1] and all of the data has been made available (see “Data availability statement”). fries/*EWAK*\* in combination achieved a significant runtime improvement over the previous state-of-the-art, *BBK*\*, with runtimes up to 2 orders of magnitude faster. *EWAK*\* also limits the number of minimized conformations used in each *K*\* score approximation by up to about 2 orders of magnitude while maintaining provable guarantees (see the Section entitled “Energy Window Approximation to K* (*EWAK*\*)”). fries alone is capable of reducing the input sequence space while provably keeping all of the most energetically favorable sequences (see the Section entitled “Fast Removal of Inadequately Energied Sequences (fries)”), decreasing the size of the sequence space by more than 2 orders of magnitude, and leading to more efficient design given the smaller search space.

To further validate osprey with fries/*EWAK*\*, we applied these algorithms to a biomedically significant design problem: the c-Raf-RBD:KRas PPI. First, we performed a series of retrospective designs where fries/*EWAK*\* accurately predicted how a variety of mutations affect the binding of c-Raf-RBD to KRas^GTP^ that previous computational methods had failed to accurately predict [49]. This success supports the use of osprey and fries/*EWAK*\* to evaluate the effect mutations in the protein-protein interface of c-Raf-RBD:KRas have on binding (more, similar successes of the *K*\* algorithm are presented and discussed in [1]). fries/*EWAK*\* also prospectively predicted the effect of new mutations in the c-Raf-RBD:KRas PPI and discovered a novel c-Raf-RBD mutation V88Y with improved affinity for KRas. We went on to combine this new mutation with two previously reported mutations, N71R and A85K [49], to create c-Raf-RBD(RKY), an even stronger binding c-Raf-RBD variant, which fries/*EWAK*\* accurately predicted. We biochemically screened top predicted variants using an initial bio-layer interferometry (BLI) single-concentration assay. Only a promising subset of the computationally predicted and initially screened variants were then evaluated using a BLI titration assay to calculate *K*_*d*_ values for individual c-Raf-RBD variants. We determined that c-Raf-RBD(RKY) binds to KRas^GTP^ roughly 36 times more tightly than wild-type c-Raf-RBD, making it the tightest known c-Raf-RBD variant binding partner of KRas^GTP^.

Given that numerous groups have explored this protein-protein interaction [64, 65, 67–77] and performed mutagenesis on c-Raf-RBD either, through rational means [64, 67, 74, 78], computational methods [49, 65] or high-throughput evolutionary methods [73, 79] and that none identified V88Y, this discovery validates our computational approach and the use of computational algorithms such as fries and *EWAK*\* to redesign protein-protein interfaces toward improved binding. Finally, previous mutations that enhanced the affinity of c-Raf-RBD binding to KRas did so by supercharging c-Raf-RBD [49, 64, 65]. In contrast, our mutation V88Y introduces a novel, aromatic residue. The discovery that such a mutation can improve the binding of c-Raf-RBD to KRas^GTP^ is of considerable significance. These new c-Raf-RBD variants serve as an important step toward better understanding the KRas:effector interface and eventually developing successful therapeutics to directly target and block the aberrant behavior of mutant KRas.

## Supporting information

**S1 Text. Homology model of c-Raf-RBD in complex with KRas.**

**S2 Text. Details of the expression and purification of c-Raf-RBD variants. S3 Text. Details of the expression and purification of KRas.**

**S1 Table. Protein structures used in computational experiments as described in the Section entitled “Computational materials and methods.”** Each protein structure has its PDB ID listed along with its molecule names as presented in the Protein Database entry for each structure. Individual designs are not listed or described here, but the necessary code and data is provided for the interested reader (see Data availability statement).

**S2 Table. Experimental and computational percent change in binding and rankings.** For each listed variant, we give the experimental percent change in binding relative to wild-type as reported in [64] and as calculated from reported binding values in [65] and [49], the *EWAK*\* computationally predicted percent change in binding (as described in the Section entitled “fries/*EWAK*\* retrospectively predicted the effect mutations in c-Raf-RBD have on binding to KRas”) and the rankings that correspond to these values. The rankings have a Pearson correlation of 0.81.

**S3 Table. Table of computational predictions for point mutants in c-R af-RBD.** Each section of the table shows the results of the redesign of a residue position in c-Raf-RBD in the c-Raf-RBD:KRas PPI in order of increasing upper bound on log(*K*\*). The table contains the values for upper and lower bounds on log(*K*\*) values (these bounds are described in detail in [32]). ^*^Design results for the wild-type amino acid identity for each position. ^*†*^Mutations that were selected for experimental testing and validation.

**S4 Table.** *K*_*d*_ **values for each tested variant for all replicates of BLI titration experiments.** For each listed variant, we give the dissociation constant *K*_*d*_ for each BLI titration experiment calculated from the fit done using the built-in mass transport model within the Octet Data Analysis HT software provided by FortéBio. We only accepted fits with a sum of square deviations *χ*^2^ less than 1 (FortéBio recommends a value less than 3) and a coefficient of determination *R*^2^ greater than 0.98. Presented in the table in Fig 10 are averages of these *K*_*d*_ values.

## Acknowledgments

We thank Rachel Kimbrough, Chanelle Simmons, Catherine Ehrhart, and Kelly Huynh for assistance with experiments and biochemical validation, Ben Fenton and Michael Kennedy for the initial KRas plasmids, and Paul Modrich for the Rosetta 2(DE3) cell lines. We also thank Terrence Oas and all members of the lab for helpful discussions.

